# Dietary adaptation of *FADS* genes in Europe varied across time and geography

**DOI:** 10.1101/111229

**Authors:** Kaixiong Ye, Feng Gao, David Wang, Ofer Bar-Yosef, Alon Keinan

**Affiliations:** Department of Biological Statistics and Computational Biology, Cornell University, Ithaca, NY, USA; Department of Anthropology, Harvard University, Cambridge, MA, USA

## Abstract

Fatty acid desaturase (*FADS*) genes encode rate-limiting enzymes for the biosynthesis of omega-6 and omega-3 long chain polyunsaturated fatty acids (LCPUFAs). This biosynthesis is essential for individuals subsisting on LCPUFAs-poor, plant-based diets. Positive selection on *FADS* genes has been reported in multiple populations, but its presence and pattern in Europeans remain elusive. Here, with analyses of ancient and modern DNA, we demonstrated that positive selection acted on the same *FADS* variants both before and after the advent of farming in Europe, but on opposite alleles. Selection in recent farmers also varied geographically, with the strongest signal in Southern Europe. These varying selection patterns concur with anthropological evidence of differences in diets, and with the association of recently-adaptive alleles with higher *FADS1* expression and enhanced LCPUFAs biosynthesis. Genome-wide association studies revealed associations of recently-adaptive alleles with not only LCPUFAs, but also other lipids and decreased risk of several inflammation-related diseases.

---

Identifying genetic adaptations to local environment, including historical dietary practice, and elucidating their implications on human health and disease are of central interest in human evolutionary genomics^1^. The fatty acid desaturase (*FADS*) gene family consists of *FADS1, FADS2* and *FADS3,* which evolved by gene duplication^2^. *FADS1* and *FADS2* encode rate-limiting enzymes for the biosynthesis of omega-3 and omega-6 long-chain polyunsaturated fatty acids (LCPUFAs) from plant-sourced shorter-chain precursors (Supplementary Fig. 1). LCPUFAs are indispensable for proper human brain development, cognitive function and immune response^3,4^. While omega-3 and omega-6 LCPUFAs can be obtained from animal-based diets, their endogenous synthesis is essential to compensate for their absence from plant-based diets. Positive selection on the *FADS* locus, a 100 kilobase (kb) region containing all three genes (Supplementary Fig. 2), has been identified in multiple populations^5-9^. Our recent study showed that a 22 bp insertion-deletion polymorphism (indel, rs66698963) within *FADS2,* which is associated with *FADS1* expression^10^, has been adaptive in Africa, South Asia and parts of East Asia, possibly driven by local historical plant-based diets^8^. We further supported this hypothesis by functional association of the adaptive insertion allele with more efficient biosynthesis^8^. In Greenlandic Inuit, who have traditionally subsisted on a LCPUFAs-rich marine diet, adaptation signals were also observed on the *FADS* locus, with adaptive alleles associated with less efficient biosynthesis^9^.

In Europeans, positive selection on the *FADS* locus has only been reported recently in a study based on ancient DNA (aDNA)^11^. Evidence of positive selection from modern DNA is still lacking even though most above studies also performed similarly-powered tests in Europeans^5-8^. Moreover, although there are well-established differences in the Neolithization process and in dietary patterns across Europe^12-14^, geographical differences of selection within Europe have not been investigated before. Furthermore, before the advent of farming, pre-Neolithic hunter-gatherers throughout Europe had been subsisting on animal-based diets with significant aquatic contribution^15-17^, in contrast to the plant-heavy diets of recent European farmers^18-20^. We hypothesized that these drastic differences in subsistence strategy and dietary practice before and after the Neolithic revolution within Europe exert different selection pressures on the *FADS* locus. In this study, we combined analyses on ancient and modern DNA to investigate potential positive selection on the *FADS* locus in Europe and to examine whether it exhibits geographical and temporal differences as would be expected from differences in diets. Briefly, we present evidence for positive selection on *opposite alleles* of the same variants before and after the Neolithic revolution, and for varying selection signals between Northern and Southern Europeans in recent history. We interpreted the functional significance of adaptive alleles with analysis of expression quantitative trait loci (eQTLs) and genome-wide association studies (GWAS), both pointing to selection for diminishing LCPUFAs biosynthesis in pre-Neolithic hunter-gatherers but for increasing biosynthesis in recent farmers. Anthropological findings indicate that these selection patterns were likely driven by dietary practice and its changes.

## Results

### Evidence of recent positive selection in Europe from both ancient and modern DNA

To systematically evaluate the presence of recent positive selection on the *FADS* locus in Europe, we performed an array of selection tests using both ancient and modern samples. We first generated a uniform set of variants across the locus in a variety of aDNA data sets (Supplementary Table S1) via imputation (Methods). For all these variants, we conducted an aDNA-based test^11^. This test includes three groups of ancient European samples and four groups of modern samples. The three ancient groups represent the three major ancestry sources of most present-day Europeans: Western and Scandinavian hunter-gatherers (WSHG), early European farmers (EF), and Steppe-Ancestry pastoralists (SA). The four groups of modern samples were drawn from the 1000 Genomes Project (1000GP), representing Tuscans (TSI), Iberians (IBS), British (GBR) and additional northern Europeans (CEU). The test identifies variants with extreme frequency change between ancient and modern samples, suggesting the presence of positive selection during recent European history (not more ancient than 8,500 years ago (ya))^11^. Our results confirmed the presence of significant selection signals on many variants in the *FADS* locus (Fig. 1), including the previously identified peak SNP rs174546 (*p* = 1.04e-21)^11^. We observed the most significant signal at an imputed SNP, rs174594 (*p* = 1.29e-24), which was not included in the original study^11^. SNP rs174570, one of the top adaptive SNPs reported in Greenlandic Inuit^9^, also exhibits a significant signal (*p* = 7.64e-18) while indel rs66698963 shows no evidence of positive selection (*p* = 3.62e-3, likely due to data quality, see Supplementary Notes). Overall, the entire peak of selection signals coincides with a linkage disequilibrium (LD) block (referred to as the *FADS1-FADS2* LD block) in Europeans, which extends over a long genomic region of 85 kb, covering the entirety of *FADS1* and most of the much longer *FADS2* (Supplementary Figs. 2 and 3). The dominant haplotype of this LD block (haplotype D; Methods) has a frequency of 63% in modern Europeans and is composed of alleles under positive selection as revealed by the above test. Of note, some alleles on this haplotype are derived (i.e. the new mutation relative to primates) while others are ancestral (Supplementary Fig. 4). Thus, the large number of variants showing genome-wide significant signals could potentially be the result of one or a few variants targeted by strong selection, with extensive hitchhiking of nearby neutral variants.

**Fig. 1.**
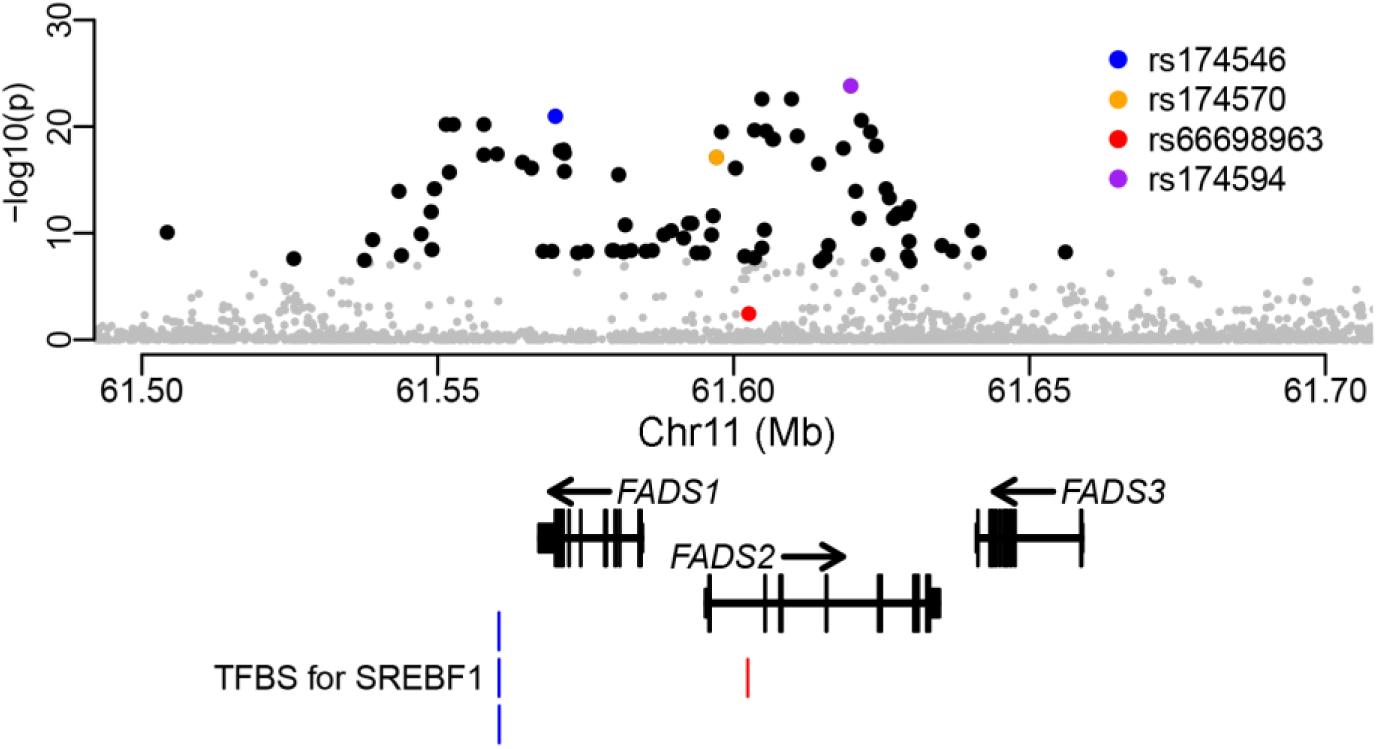
Ancient DNA-based test for recent positive selection. The y axis indicates genomic control corrected *p* values at a negative logarithm scale. Variants under genome-wide significance level (5e-8) are in gray except for highlighted ones. Four variants are highlighted: the most significant SNP (purple); the top SNP reported by Mathieson *etal.^11^* (blue); one of the top adaptive SNPs reported in Greenlandic Inuit^9^ (orange); the indel reported to be targeted by positive selection in populations with historical plant-based diets^8^ (red). The overall pattern is consistent with that previously described^11^ (Supplementary Fig. S31). At the bottom are the representative transcript models for the three *FADS* genes and the four transcription factor binding sites (TFBS) for SREBF1 from ENCODE^84^ (blue) and another previous study^10^ (red).

We next performed several selection tests solely based on extant European populations. Considering the five European populations from 1000GP, including samples of Finns (FIN) and the four samples described above, a haplotype-based selection test, nSL^21^, revealed positive selection on many SNPs in the *FADS1-FADS2* LD block. Importantly, this test unraveled the same adaptive alleles as in the above aDNA-based test and a same general trend of stronger signal towards rs174594 (Fig. 2A, Supplementary Fig. 5). For rs174594, the nSL values are significant in all five populations and the signal exhibits a gradient of being stronger in southern Europeans and weaker in northern Europeans (Fig. 2A, Supplementary Fig. 6): TSI (*p* = 0.00044), IBS (*p* = 0.0020), CEU (*p* = 0.0039), GBR (*p* = 0.0093), and FIN (*p* = 0.017). Of note, nSL values have been normalized separately in each population to remove demographic effects^21^. The other three variants of interest (rs174546, rs174570, and rs66698963) exhibit no selection signals, except for rs174570 showing borderline significance in the two southernmost populations (TSI: *p* = 0.022; IBS: *p* = 0.050, Fig. 2A). Signals were also observed with nSL in two whole-genome sequencing cohorts of British ancestry from the UK10K project (Supplementary Fig. 7). Another test for positive selection in very recent history (during the past ~2,000-3,000 years), Singleton Density Score (SDS)^22^, applied in the UK10K data set, also revealed significant signals for multiple SNPs in the *FADS1-FADS2* LD block, with the same adaptive alleles and general trend of localized signals as in the above two tests (Fig. 2B, Supplementary Fig. S8). Significant SDS was observed for rs174594 (*p* = 0.045) and rs174570 (*p* = 0.045), but not rs174546. Of note, it is the derived allele for rs174594 that was under selection, while it is the ancestral allele for rs174570. Interestingly, selection on the opposite (or derived) allele of rs174570 has been shown in Greenlandic Inuit^9^ Additional tests of selection consistently highlight the *FADS1-FADS2* LD block as a target of natural selection (Supplementary Figs. S5, S7-S10). Taken together, standard tests on modern DNAs support the aDNA-based results of recent positive selection on the *FADS* locus and, specifically, on the D haplotype of the *FADS1-FADS2* LD block.

**Fig. 2.**
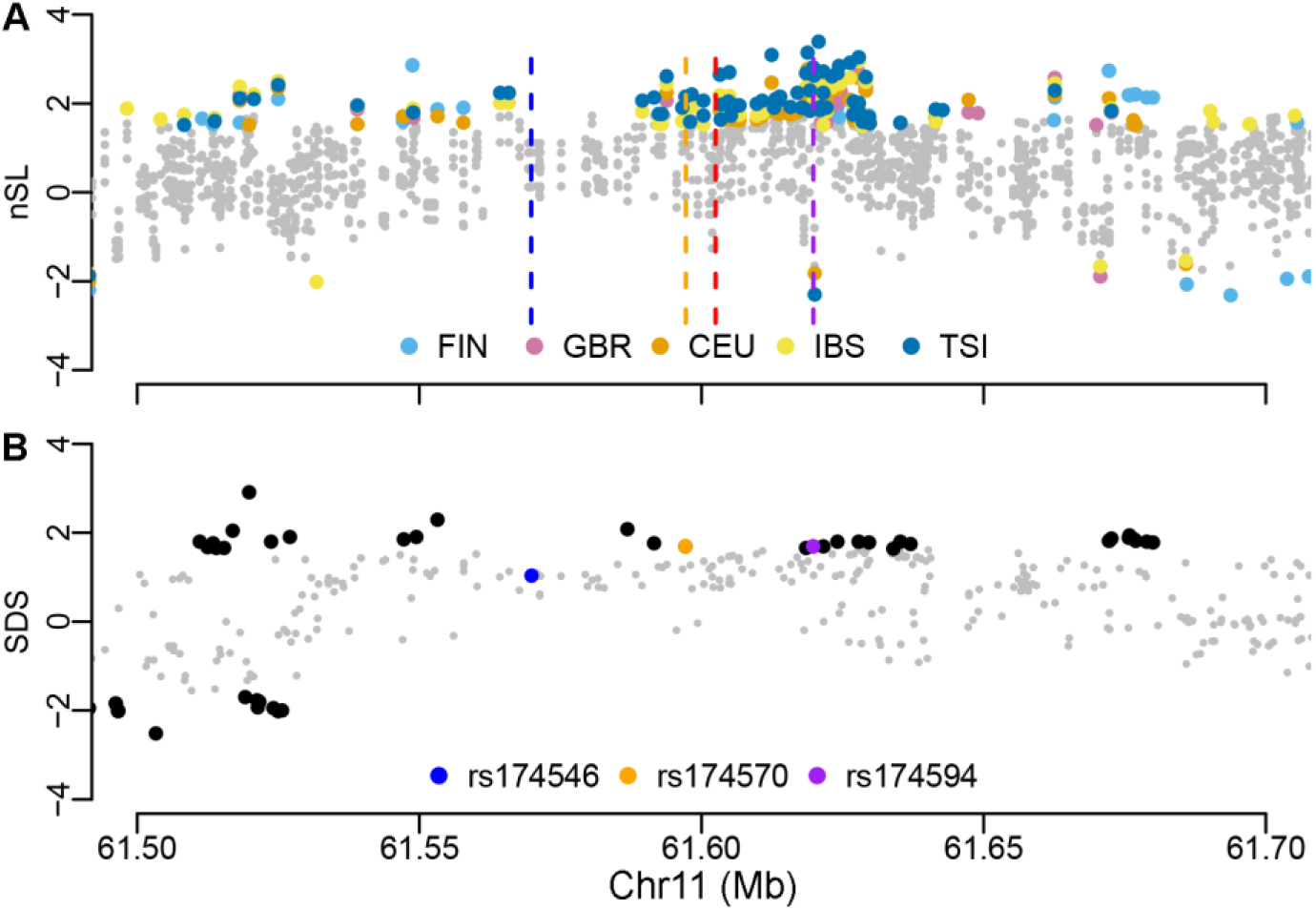
Tests for recent positive selection based solely on modern DNA. (**A**) Haplotype-based selection test (nSL^21^) in modern Europeans from 1000GP. The test was performed separately for each of the five European groups. Only variants with significant values are shown with population-specific colors as indicated in the legend. The positions for four variants of interest were indicated with vertical dashed lines, colored as in Fig. 1. For presentation purpose, the sign was set so that being positive indicates that the adaptive allele revealed by nSL is consistent with that revealed by the aDNA-based test in Fig. 1. Original statistics for 1000GP and UK10K are shown in Supplementary Figs. S5 and S7. The five 1000GP European populations are: CEU - Utah Residents (CEPH) with Northern and Western Ancestry; FIN - Finnish in Finland; GRB - British in England and Scotland; IBS - Iberian Population in Spain; TSI - Toscani in Italia. (**B**) Singleton Density Score (SDS^22^) in modern Europeans from UK10K. Variants under significance level are in gray except for highlighted ones. Three variants of interest were highlighted with colors as indicated in the legend. The indel rs66698963 was not present in the original UK10K data set. The sign of SDS was set as in nSL. Original statistics are shown in Supplementary Fig. S8.

### Geographical differences of recent positive selection signals across Europe

To rigorously evaluate geographical differences of recent positive selection on the *FADS* locus across Europe, we revisited the aDNA-based selection test^11^. We started by decomposing the original test for four representative SNPs (Fig. 3A) and then performed the test separately in Northern and Southern Europeans for all variants in the *FADS* locus (Fig. 3B). Our first analysis included four SNPs, three of which (rs174594, rs174546, and rs174570) are top SNPs from this and previous studies^9,11^ and are highlighted in all our analyses, while the fourth (rs4246215) is the one showing the biggest difference in the upcoming South-North comparison analysis. The indel rs66698963 was not highlighted in this and all upcoming analyses because it has no significant selection signals in Europe. The original aDNA-based test evaluates the frequencies of an allele in three ancient samples and four modern 1000GP samples under two hypotheses (H_0_ and H_1_). Under H_1_, maximum likelihood estimates (MLEs) of frequencies in all samples are constrained only by observed allele counts and thus equivalent to the direct observed frequencies (Fig. 3A; blue bars). Among the four modern samples, the observed adaptive allele frequencies for all four SNPs exhibit a South-North gradient with the highest in Tuscans and the lowest in Finns, consistent with the gradient of selection signals observed before based on modern DNA. Among the three ancient samples, the observed allele frequencies, equivalent to the frequencies upon admixture (Fig. 3A, orange bars for ancient groups), are always the lowest and often zero in the WSHG sample.

**Fig. 3.**
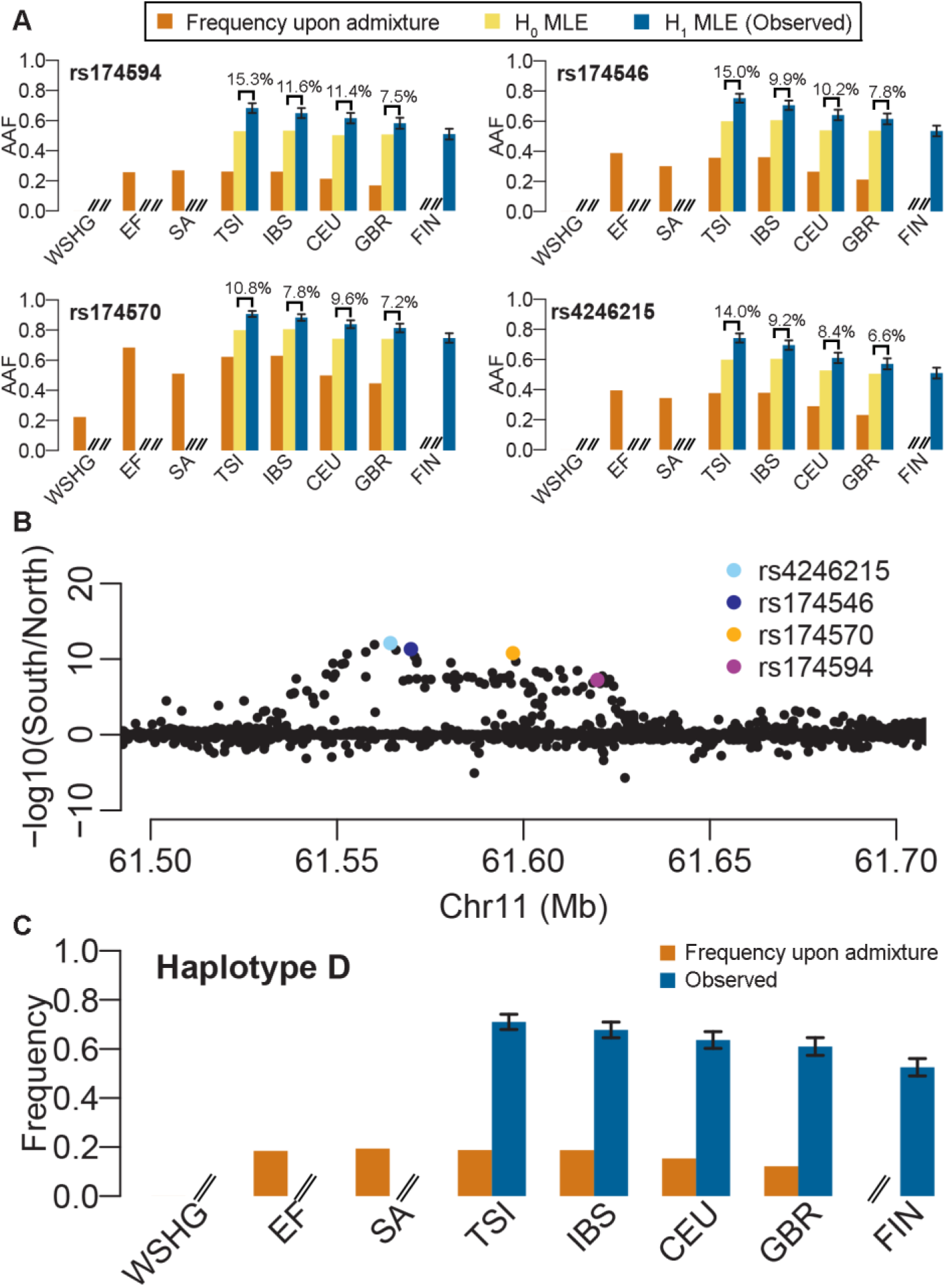
Varying selection and frequency patterns between Southern and Northern Europe. (**A**) South-North frequency gradient for adaptive alleles of four representative SNPs under different scenarios. AAF refers to adaptive allele frequency. Orange bars represent frequencies upon admixture, which were directly observed in ancient groups and predicted for extant populations based on linear mixture of frequencies in ancient groups. Yellow bars represent frequencies estimated under H_0_. Estimates for ancient groups were not shown because they are not relevant here. Blue bars represent frequencies estimated under H_1_, whose only constraint is the observed data and therefore the MLEs are just the observed means. The estimates for ancient groups are the same as their frequencies upon admixture and are omitted on the plot. The absolute difference between H0 and H1 estimates are indicated above the corresponding bars. Please note that the frequencies upon admixture in WSHG are 0 for rs174594, rs174546 and rs4246215 and no bars were plotted. (**B**) Comparison of aDNA-based selection signals between Southern and Northern Europe. aDNA-based selection tests were performed separately for Southern (TSI and IBS) and Northern (CEU and GBR) Europeans. For each variant, the *p* values from these two tests were compared at a −log10 scale (y axis). SNPs of interest were colored as indicated. (**C**) South-North frequency gradient for the adaptive haplotype in extant populations. The two frequency types are just as in (A). The frequency upon admixture for WSHG is 0. In (A) and (C), FIN has only observed values. If values are not shown or not available, signs of “//” are indicated at corresponding positions. Error bars stand for standard errors.

Under H_0_, the MLEs of frequencies are constrained by the observed allele counts and an additional assumption that an allele’s frequencies in the four modern samples are each a linear combination of its frequencies in the three ancient samples. Considering the later assumption alone, we can predict the frequencies of adaptive alleles right after admixture for each modern population. The admixture contribution of WSHG, as estimated genome-wide, is higher towards the North, constituting of 0%, 0%, 19.6%, and 36.2% for TSI, IBS, CEU, and GBR, respectively^11^. Thus, the predicted adaptive allele frequencies upon admixture for these four modern populations are usually lower in the North (Fig. 3A; orange bars in modern populations), suggesting higher starting frequencies in the South at the onset of selection. Further considering observed allele counts, we obtained the MLEs of frequencies under H_0_ (Fig. 3A; yellow bars in modern populations). As expected, the predicted allele frequencies are higher in the South. But more importantly, the differences between H0 and H1 estimates in modern populations (Fig. 3A; indicated differences between yellow and blue bars) are still higher in the South, suggesting that in addition to population-specific admixture proportions and different starting frequencies, more recent factors, such as stronger selection pressure, earlier onset of selection, or unmodeled recent demographic history, might contribute to the observation of stronger selection signals in the South.

To evaluate the potential confounding effects of varying demographic history that is not captured by the model, we evaluate all variants in the 3 Mb region surrounding the *FADS* locus. We applied the aDNA-based selection test separately for the two Southern and the two Northern populations. All variants that were significant in the combined analyses (Fig. 1) were also significant in each of the two separate analyses, but many exhibited much stronger signals in Southern populations (Fig. 3B; Supplementary Fig. S11). The maximum difference was found for SNP rs4246215, whose *p* value in Southern populations is 12 orders of magnitude stronger than that in Northern populations. SNP rs174594, rs174546 and rs174570 also have signals that are several orders of magnitude stronger in the South. A further decomposition of the selection test and comparison of maximum likelihoods under H_0_ and H_1_ between South and North revealed that a stronger deviation under H_0_ in the South is driving the signal (Supplementary Fig. S12). It is noteworthy that the pattern of stronger signal in the South is observed only for some but not all SNPs, excluding the possibility of systemic bias and pointing at variants-specific properties, likely for variants that were under selection and the nearby variants in LD. Indeed, the candidate adaptive haplotype D also exhibits frequency patterns that are consistent with adaptive alleles of the four representative SNPs (Fig. 3C). Hence, these results suggest that there might be stronger selection pressure or earlier onset of positive selection on the *FADS1-FADS2* LD block in Southern Europeans.

### Opposite selection signals in pre-Neolithic European hunter-gatherers

Motivated by the very different diet of pre-Neolithic European hunter-gatherers, we set to test the action of natural selection on the *FADS* locus before the Neolithic revolution. We started by examining the frequency trajectory of haplotype D, the candidate adaptive haplotype in recent European history. As noted above, its frequency increased drastically in Europe after the Neolithic revolution (Fig. 3C, the contrast between orange and blue bars). In stark contrast, it shows a clear trajectory of decreasing frequency over time among pre-Neolithic hunter-gatherers^23^ (Fig. 4A): starting from 32% in the ∼30,000-year-old (yo) “Věstonice cluster”, through 21% in the ∼ 15,000 yo “El Mirón cluster”, to 13% in the ∼ 10,000 yo “Villabruna cluster”, and to being practically absent in the ∼ 7,500 yo WSHG group. We hypothesized that there was positive selection on alleles opposite to the recently adaptive ones on haplotype D.

**Fig. 4.**
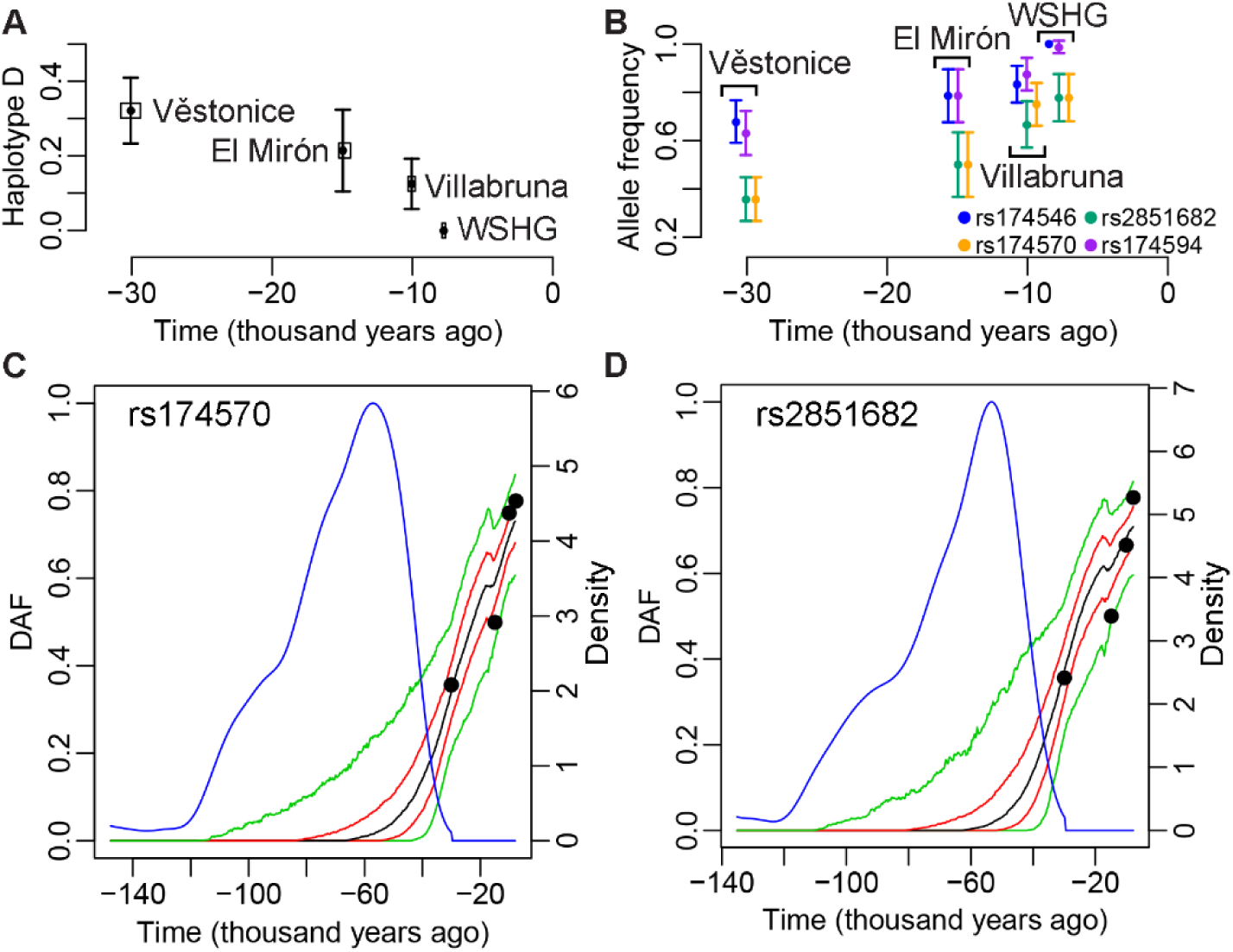
Temporal frequency pattern and selection signals in pre-Neolithic European hunter-gatherers. (**A**) The frequency of haplotype D over time in four groups of hunter-gatherers. Frequency for each group is plotted as a black point at the median age of samples. The horizontal box surrounding the point represents the medians of lower- and upper-bound estimates of sample ages. Error bars are standard errors. Group names are indicated next to their frequencies. The frequency for WSHG is 0. (**B**) Allele frequencies for four SNPs. It has similar format as in (A) except that small arbitrary values were added on their x coordinates in order to visualize all SNPs, which were colored as indicated in the legend. The alleles chosen are the ones increasing frequency over time. They are derived alleles for rs174570 and rs2851682, and ancestral alleles for rs174546 and rs174594. (**C**) and (**D**) Posterior distribution on the derived allele frequency path for rs174570 and rs2851682, respectively. The sampled frequencies are indicated with black points, which are the same point estimates as in (B). The median, 25% and 75% quantiles, and 5% and 95% quantiles of the posterior distribution are indicated respectively with black, red and green lines. The posterior distribution on the age of derived allele is shown with a blue line, with values on the right y axis.

To search for variants with evidence of positive selection during the pre-Neolithic period, we considered the allele frequency time series for all variants around the *FADS* locus. We applied to each variant two rigorous, recently-published Bayesian methods^24,25^ to infer selection coefficients from time series data. Under a simple demographic model of constant population size, both methods consistently highlighted two SNPs (rs174570 and rs2851682) within the *FADS1-FADS2* LD block to be under positive selection during the pre-Neolithic period tested, approximately 30,000-7,500 ya (Supplementary Figs. 13 and 14). The Schraiber *et al.* method is capable of processing more complicated demographic models^24^. With this method and considering a more realistic European demographic model^26^, the same two SNPs were highlighted (Supplementary Fig. 15). The derived alleles of these two SNPs have similar frequency trajectories during the examined period, increasing from 36% to 78% (Fig. 4B).

Estimated selection coefficients for homozygotes of adaptive allele (s) for these two SNPs are similar across methods and demographic models. With the Schraiber *et al.* method and the realistic demographic model, the marginal maximum *a posteriori* estimate of s for rs174570 is 0.38% (95% credible interval (CI): 0.038% – 0.92%) while the estimated derived allele age is 57,380 years (95% CI: 157,690 – 41,930 years) (Fig. 4C, Supplementary Fig. 16). For rs2851682, the estimated s is 0.40% (95% CI: 0.028% – 1.12%) while its derived allele age is 53,440 years (95% CI: 139,620 – 39,320 years) (Fig. 4D, Supplementary Fig. 17). In addition to these two SNPs, ApproxWF^25^ revealed significant signals for 44 SNPs in the *FADS1-FADS2* LD block (Supplementary Fig. 14), including rs174546 and rs174594, whose ancestral allele frequencies increased from about 65% to almost fixation (Fig. 4B). Importantly, these SNPs have similar estimated s (0.28% - 0.62%) and their adaptive alleles are alternative (or opposite) to the ones under selection in recent history.

Considering the haplotype structure of the *FADS1-FADS2* LD block (Fig. 5A), we identified a haplotype (referred to as M2), which is comprised of alleles that are mostly alternative to those on haplotype D (Supplementary Fig. S4). M2 appears in modern Europeans at a frequency of 10% but is much more common in Eskimos from Eastern Siberia, presumably for similar reasons that the derived allele of rs174570 is prevalent in Greenlandic Inuit. M2 exhibits increasing frequency over time in pre-Neolithic hunter-gatherers (Supplementary Table S2), suggesting that the allele(s) targeted by selection during that period are likely on M2.

**Fig. 5.**
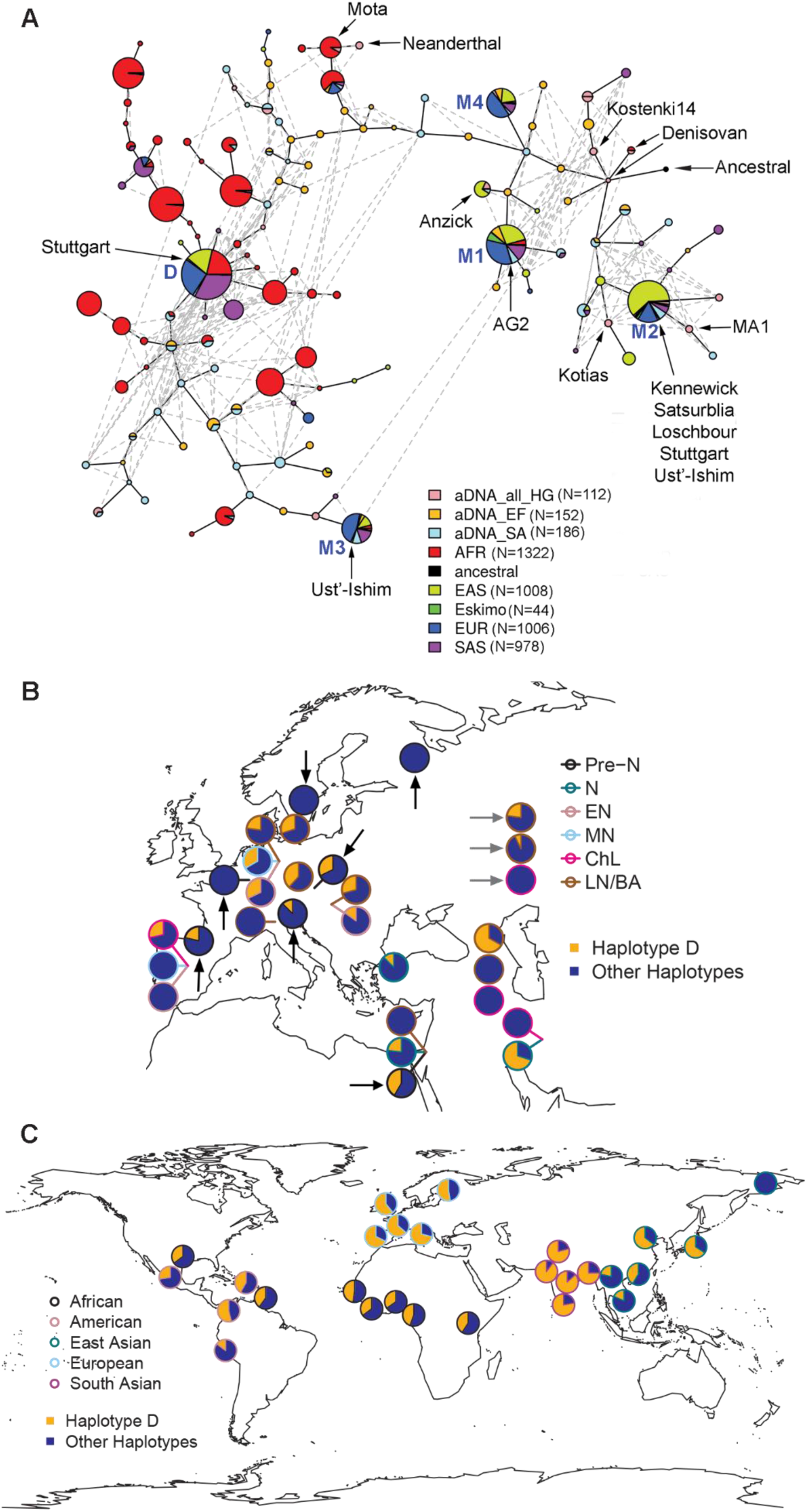
Haplotype network and geographical frequency distribution. (**A**) Haplotype network for 1000G samples (2,157 individuals, excluding admixed American samples), 22 modern Eskimos and 225 aDNAs. Each pie chart represents one haplotype and its size is proportional to log_2_(# of haplotype) plus a minimum size to visualize rare haplotypes. Sections in the pie provide the breakdown by groups. Detailed haplotype frequencies are in Supplementary Table S2. The edges connecting haplotypes are of arbitrary length. Haplotypes for some well-known ancient samples are labelled. The top five haplotypes in modern Europeans, referred to as D, M1, M2, M3, and M4 from the most to least frequent common, are indicated with their names in blue. (**B**) Frequency of haplotype D in Eurasian ancient DNAs. Each pie represents one sampled group and is placed at the sampling location or nearby with a line pointing at the sampling location. The color of the pie chart border indicates the archaeological period. If multiple samples of different periods were collected at the same geographical location, these samples are ordered vertically with the older samples at the bottom. Hunter-gatherer groups are indicated with black arrows and pastoralist groups with gray arrows, while others are farmers. Geographical locations for some hunter-gatherer groups (e.g. the Věstonice, El Mirón and Villabruna clusters) are only from representative samples. Detailed frequencies are in Supplementary Table S3. Pre-N: Pre-Neolithic; N: Neolithic; EN: Early Neolithic; MN: Mid-Neolithic; ChL: Chalcolithic; LN/BA: Late Neolithic/Bronze Age. (**C**) Frequency of haplotype D in present-day global populations. All 26 populations from 1000GP and one Eskimo group are included. The color of the pie chart border represents the genetic ancestry. It is noteworthy that there are two samples in America that are actually of African ancestry. Detailed frequencies are in Supplementary Table S4.

### The temporal and global evolutionary trajectory of *FADS* haplotypes

To go beyond the dominant D haplotype and study different haplotypes in the *FADS1-FADS2* LD block, their frequency changes over time and their current global distributions, we performed haplotype network and frequency analysis on 450 and 5,052 haplotypes from ancient and modern DNA, respectively (Fig. 5, Supplementary Fig. S18, Supplementary Tables S2-S4). The top five haplotypes in modern Europeans, designated as D, M1, M2, M3 and M4 from the most to the least common (63%, 15%, 10%, 5%, 4%, respectively), were all present in aDNA and modern Africans. M1, M2 and M4 are closer to the consensus ancestral haplotype observed in primates while D and M3 are more distant (Fig. 5A). Among the Out-of-Africa ancestors, the frequencies of D and M2 were probably ∼35% and ∼27% because those frequencies were observed in both the oldest European hunter-gatherer group, the ∼30,000 yo “Věstonice cluster”, and the ∼14,500 yo Epipalaeolithic Natufian hunter-gatherers in the Levant (Fig. 5B, Supplementary Table S3). Among pre-Neolithic European hunter-gatherers, positive selection on M2 increased its frequency from 29% to 56% from approximately 30,000 to 7,500 ya, while the D haplotype practically disappeared by the advent of farming (Figs. 4A and 5B). With the arrival of farmers and Steppe-Ancestry pastoralists, D was re-introduced into Europe. Since the Neolithic revolution, positive selection on D increased its frequency dramatically to 63% while that of M2 has decreased to only 10% among present-day Europeans. Globally, D is also present at high frequency in South Asia (82%) but absent in modern-day Eskimos (Fig. 5C). In contract, M2 has very low frequency in South Asia (3%) but moderate frequency in Eskimos (27%). Further detailed description of evolutionary trajectories of haplotypes in this region could be found in Supplementary Notes.

The geographical frequency patterns of representative variants (rs174570, rs174594, rs174546, rs66698963, and rs2851682; Fig. 6, Supplementary Figs. 19-23) mostly mirror those of key haplotypes, but with discrepancies providing insights into casual variants and allele ages. One major discrepancy was found in Africa. The derived alleles of rs174570 and rs2851682 remain almost absent in Africa, consistent with their allele age estimates of ~55,000 years (Figs. 4C and 4D) and ruling out their involvement in the positive selection on *FADS* genes in Africa^5,6,8^. Considering the much weaker LD structure of the *FADS* locus in Africa (Supplementary Fig.24), it is possible that selection in Africa may be on haplotypes and variants that are different from those in Europe.

**Fig. 6.**
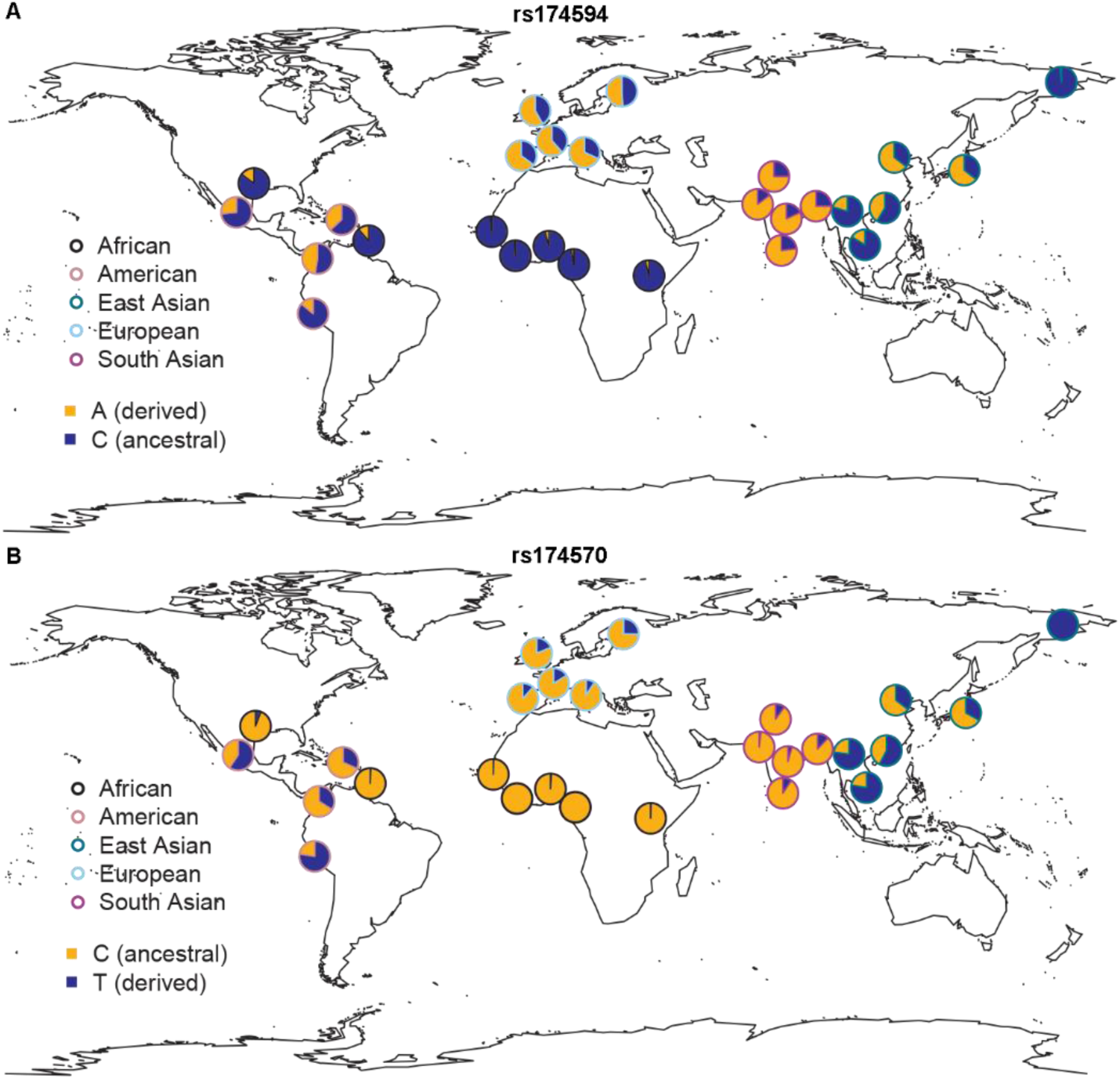
Geographical frequency distribution for SNPs rs174594 and rs174570 in present-day global populations. Adaptive alleles in recent European history are colored in orange. All 26 populations from 1000GP and one Eskimo group are included. The color of the pie chart border represents the genetic ancestry. It is noteworthy that there are two samples in America that are actually of African ancestry. Similar global patterns were observed with HGDP samples (Supplementary Figs. S19 and S21).

### Functional and medical implications of adaptive variants

Previous studies on adaptive evolution of the *FADS* locus suggested that adaptive alleles are associated with expression levels of *FADS* genes^5,6,8^. To test this possibility in the context of this large-sale analysis, we considered data from the Genotype-Tissue Expression (GTEx) project^27^. Our results point to many SNPs on the *FADS1-FADS2* LD block being eQTLs of *FADS* genes. Out of a total of 44 tissues, these eQTLs at genome-wide significance level are associated with the expression of *FADS1, FADS2,* and *FADS3* in 12, 23, and 4 tissues, respectively, for a total of 27 tissues (Supplementary Figs. 25-27). Considering the peak SNP rs174594 alone, nominally significant associations with these three genes were found in 29, 28 and 4 tissues, respectively. Importantly, out of these tissues with association signals, the adaptive allele in recent European history is associated with higher expression of *FADS1,* lower expression of *FADS2* and higher expression of *FADS3* in 28, 27 and 4 tissues, respectively. The general trend that recently adaptive allele is associated with higher expression of *FADS1* but lower expression of *FADS2* was also observed for other representative SNPs (rs174546, rs174570, and rs2851682).

GWAS have revealed 178 association signals with 44 different traits in the *FADS1-FADS2* LD block, as recorded in the GWAS catalog (Supplementary Tables S5-S9)^28^. All effects reported in the following are based on individuals of European ancestry, while some are also replicated in other ethnic groups. We report the direction of association in terms of recently adaptive alleles, while the direction is opposite for adaptive alleles in pre-Neolithic hunter-gatherers. Dissecting different associations, (1) the most prominent group of associated traits are polyunsaturated fatty acids (PUFAs, Supplementary Fig. S1), including LCPUFAs and their shorter-chain precursors. Alleles on haplotype D are associated with higher levels of arachidonic acid (AA)^29-31^, adrenic acid (AdrA)^29,31-33^, eicosapentaenoic acid (EPA)^31,34^ and docosapentaenoic acid (DPA)^31,32,34^, but with lower levels of dihomo-gamma-linolenic acid (DGLA)^29-32^, all of which suggest increased activity of delta-5 desaturase encoded by FADS1^31,35^. This is consistent with the association of recently adaptive alleles with higher *FADS1* expression. Surprisingly, these alleles are associated with higher levels of gamma-linolenic acid (GLA)^29,30,32^ and stearidonic acid (SDA)^31^, but with lower levels of linoleic acid (LA)^29,30,32,36^ and alpha-linolenic acid (ALA)^30,32,34^, suggesting increased activity of delta-6 desaturase encoded by *FADS2*^30^. However, the above eQTL analysis suggested that recently adaptive alleles tend to be associated with lower *FADS2* expression. Some of these association signals have been replicated across Europeans^29,31-36^, Africans^34^, East Asians^30,34^, and Hispanic/Latino^34^. (2) Besides PUFAs, recently adaptive alleles are associated with decreased cis/trans-18:2 fatty acids^37^, which in turn is associated with lower risks for systemic inflammation and cardiac death^37^. Consistently, these alleles are also associated with decreased resting heart rate^38,39^, which reduces risks of cardiovascular disease and mortality. (3) With regards to other lipid levels, recently adaptive alleles have been associated with higher levels of high-density lipoprotein cholesterol (HDL)^40-45^, low-density lipoprotein cholesterol (LDL)^40-42,46^ and total cholesterol^40-42^, but with lower levels of triglycerides^40,41,44,45^. (4) In terms of direct association with disease risk, these alleles are associated with lower risk of inflammatory bowel diseases, both Crohn’s disease^47-49^ and ulcerative colitis^49^, and of bipolar disorder^50^.

Going beyond known associations from the GWAS catalog, we analyzed data from the two sequencing cohorts of the UK10K study. Focusing on the peak SNP rs174594, we confirmed the association of the recently adaptive allele with higher levels of TC, LDL, and HDL. We further revealed its association with higher levels of additional lipids, Apo A1 and Apo B (Supplementary Fig. 28). Taken together, recently adaptive alleles, beyond their direct association with fatty acid levels, are associated with factors that are mostly protective against inflammatory and cardiovascular diseases, and indeed show direct association with decreased risk of inflammatory bowel diseases.

## Discussion

Recent positive selection on *FADS* genes after the Neolithic revolution in Europe has been previously reported^11^. Here, we provided a more detailed view of this recent selection and revealed that it varied geographically, between the North and the South (Figs. 1-3). We further discovered a unique phenomenon that before the Neolithic revolution, the same variants were also subject to positive selection, but with the opposite alleles being selected (Fig. 4). We showed that alleles diminishing LCPUFAs biosynthesis were adaptive before the Neolithic revolution, while alleles enhancing LCPUFAs biosynthesis were adaptive after the Neolithic revolution. In Supplementary Notes, we provided detailed discussions of our results, including 1) interpreting results from different selection tests, especially considering the complications of selection on alternative alleles in two historic periods and selection on standing variations in recent history; 2) interpreting results concerning South-North differences, including consideration of potential geographical differences in demographic history; 3) interpreting eQTLs and GWAS results. Here, we focus instead on interpreting the selection patterns in light of anthropological findings.

The dispersal of the Neolithic package into Europe about 8,500 ya caused a sharp dietary shift from an animal-based diet with significant aquatic contribution to a terrestrial plant-heavy diet including dairy products^15-20^. For pre-Neolithic European hunter-gatherers, the significant role of aquatic food, either marine or freshwater, has been established in sites along the Atlantic coast^17,51-53^, around the Baltic sea^17^, and along the Danube river^54^. The content of LCPUFAs are usually the highest in aquatic foods, lower in animal meat and milk, and almost negligible in most plants^55^. Consistent with the subsistence strategy and dietary pattern, positive selection on *FADS* genes in pre-Neolithic hunter-gatherers was on alleles associated with less efficient LCPUFAs biosynthesis, possibly compensating for the high dietary input. In addition to obtaining sufficient amounts of LCPUFAs, maintaining a balanced ratio of omega-6 to omega-3 is critical for human health^56^. Hence, it is also plausible that positive selection in hunter-gatherers was in response to an unbalanced omega-6 to omega-3 ratio (e.g. too much omega-3 LCPUFAs). Positive selection on *FADS* genes has also been observed in modern Greenlandic Inuit, who subsist on a seafood diet^9^. Specifically, the derived allele of rs174570 exhibits positive selection signals in both pre-Neolithic European hunter-gatherers and extant Greenlandic Inuit. More generally, haplotype M2, the candidate adaptive haplotype in pre-Neolithic Europe, is also common in the extant Eskimo samples examined in our study. It is noteworthy that aquatic food was less prevalent among pre-Neolithic hunter-gatherers around the Mediterranean basin, possibly due to the low productivity of the Mediterranean Sea^57-59^. It would be interesting to examine the geographical differences of selection in pre-Neolithic Europe. However, pre-Neolithic aDNA is still scarce, prohibiting such an analysis at present.

The Neolithization of Europe^12,60,61^ started in the Southeast region around 8,500 ya when farming and herding spread into the Aegean and the Balkans. Despite a few temporary stops, it continued spreading into central and northern Europe following the Danube River and its tributaries, and along the Mediterranean coast. It arrived at the Italian Peninsula about 8,000 ya and shortly after reached Iberia by 7,500 ya. While farming rapidly spread across the loess plains of Central Europe and reached the Paris Basin by 7,000 ya, it took another 1,000 or more years before it spread into Britain and Northern Europe around 6,000 ya. From that time on, European farmers relied heavily on their domesticated animals and plants. Compared to pre-Neolithic hunter-gatherers, European farmers consumed much more plants and less aquatic foods^18-20,62^. Consistent with the lack of LCPUFAs in plant-based diets, positive selection on *FADS* genes during recent European history has been on alleles associated with enhanced LCPUFAs biosynthesis from plant-derived precursors (LA and ALA). Positive selection for enhanced LCPUFAs synthesis has also been observed previously in Africans, South Asians and some East Asians, possibly driven by their traditional plant-based diets^5,6,8^.

Despite the overall trend of relying heavily on domesticated plants, there are geographical differences of dietary patterns among European farmers. In addition to the 2,000-year-late arrival of farming at Northern Europe, animal husbandry and the consumption of animal milk became gradually more prevalent as Neolithic farmers spread to the Northwest^18,61,63-65^. Moreover, similar to their pre-Neolithic predecessors, Northwestern European farmers close to the Atlantic Ocean or the Baltic Sea still consumed more marine food than their Southern counterparts in the Mediterranean basin^66,67^. It is noteworthy that historic dairying practice in Northwestern Europe has driven the adaptive evolution of lactase persistence in Europe to reach the highest prevalence in this region^64^. In this study, we observed that recent selection signals for alleles enhancing LCPUFAs biosynthesis are stronger in Southern than in Northern Europeans, even after considering the later arrival of farming and the lower starting allele frequencies in the North. The higher aquatic contribution and stronger reliance on animal meat and milk might be responsible for a weaker selection pressure in the North. However, since GWAS results have unraveled many traits and diseases associated with *FADS* genes, it is possible that other environmental factors beyond diet were involved.

## Conclusions

We presented several lines of evidence for positive selection on *FADS* genes in Europe and for its geographically and temporally varying patterns. These patterns concur with mounting anthropological evidence of geographical variability and historical change in dietary patterns. Specifically, in pre-Neolithic hunter-gatherers subsisting on animal-based diets with significant aquatic contribution, LCPUFAs-synthesis-diminishing alleles have been adaptive. In recent European farmers subsisting on plant-heavy diets, LCPUFAs-synthesis-enhancing alleles have been adaptive. Importantly, these are not simply any alleles with opposite functional consequence, but are alternative alleles of the same variants such that when one is under selection and increases in frequency, the other will decrease in frequency. To the best of our knowledge, this is the first example of its kind in humans. Moreover, we reported geographically varying patterns of recent selection that are in line with a stronger dietary reliance on plants in Southern European farmers. These unique, varying patterns of positive selection in different dietary environments, together with the large number of traits and diseases associated with the adaptive region, highlight the importance and potential of matching diet to genome in the future nutritional practice.

## Methods

### Data sets

The ancient DNA (aDNA) data set was compiled from two previous studies^23,68^, which in turn were assembled from many studies, in addition to new sequenced samples. These two data sets were merged by removing overlapping samples. In total, there are 325 ancient samples included in this study. Information about these samples and their original references could be found in Supplementary Table S1. For the aDNA-based test for recent selection, a subset of 178 ancient samples were used and clustered into three groups as in the original study^11^, representing the three major ancestral sources for most present-day European populations. These three groups are: West and Scandinavian hunter-gatherers (WSHG, N=9), early European farmers (EF, N=76), and individuals of Steppe-pastoralist Ancestry (SA, N=93). Three samples in the EF group in the original study were excluded from our analysis because they are genetic outliers to this group based on additional analysis^68^. For aDNA-based tests for ancient selection in pre-Neolithic European hunter-gatherers, a subset of 42 ancient samples were used and four groups were defined. In addition to the WSHG (N=9), the other three groups were as originally defined in a previous study^23^: the “Věstonice cluster”, composed of 14 pre-Last Glacial Maximum individuals from 34,000-26,000 ya; the “El Mirón cluster”, composed of 7 post-Last Glacial Maximum individuals from 19,000-14,000 ya; the “Villabruna cluster”, composed of 12 post-Last Glacial Maximum individuals from 14,000-7,000 ya. There were three Western hunter-gatherers that were originally included in the “Villabruna cluster”^23^, but we included them in WSHG in the current study because of their similar ages in addition to genetic affinity^11^. In haplotype network analysis, all aDNAs included in the two aDNA-based selection tests were also included. In addition, we included some well-known ancient samples, such as the Neanderthal, Denisovan, and Ust’-Ishim. In total, there were 225 ancient samples (450 haplotypes). For geographical frequency distribution analysis, a total of 300 ancient samples were used and classified into 29 previously defined groups^11,23,68^ based on their genetic affinity, sampling locations and estimated ages.

The 1000 Genomes Project (1000GP, phase 3)^7^ has sequencing-based genome-wide SNPs for 2,504 individuals from 5 continental regions and 26 global populations. Detailed description of these populations and their sample sizes are in Supplementary Methods. The Human Genome Diversity Project (HGDP)^69^ has genotyping-based genome-wide SNPs for 939 unrelated individuals from 51 populations. The data from the Population Reference Sample (POPRES)^70^ were retrieved from dbGaP with permission. Only 3,192 Europeans were included in our analysis. The country of origin of each sample was defined with two approaches. Firstly, a “strict consensus” approach was used: an individual’s country of origin was called if and only if all four of his/her grandparents shared the same country of origin. Secondly, a more inclusive approach was used to further include individuals that had no information about their grandparents. In this case, their countries of birth were used. Both approaches yielded similar results and only results from the inclusive approach are reported. The 22 Eskimo samples were extracted from the Human Origins dataset^71^.

The two sequencing cohorts of UK10K were obtained from European Genome-phenome Archive with permission^72^. These two cohorts, called ALSPAC and TwinsUK, included low-depth whole-genome sequencing data and a range of quantitative traits for 3,781 British individuals of European ancestry (N=1,927 and 1,854 for ALSPAC and TwinsUK, respectively)^72^.

### Imputation for ancient and modern DNA

Genotype imputation was performed using Beagle 4.1^73^ separately for data sets of aDNA, HGDP and POPRES. The 1000GP phase 3 data were used as the reference panel^7^. Imputation was performed for a 5-Mb region surrounding the *FADS* locus (hg19:chr11: 59,100,000-64,100,000), although most of our analysis was restricted to a 200 kb region (hg19:chr11:61,500,000-61,700,000). For most of our analysis (e.g. estimated allele count or frequency for each group), genotype probabilities were taken into account without setting a specific cutoff. For haplotype-based analysis (e.g. estimated haplotype frequency for each group), a cutoff of 0.8 was enforced and haplotypes were defined with missing data and following the phasing information from imputation.

Genotype imputation for aDNA has been shown to be desirable and reliable^74^. We also evaluated the imputation quality for aDNA by comparing with the two modern data sets (Supplementary Fig. S29). Overall, the imputation accuracy for ungenotyped SNPs, measured with allelic R^2^ and dosage R^2^, is comparable between aDNA and HGDP, but is higher in aDNA when compared with POPRES. Note that sample sizes are much larger for HGDP (N=939) and POPRES (N=3,192), compared to aDNA (N=325). The comparable or even higher imputation quality in aDNA was achieved because of the higher density of genotyped SNPs in the region.

### Linkage disequilibrium and haplotype network analysis

Linkage disequilibrium (LD) analysis was performed with the Haploview software (version 4.2)^75^. Analysis was performed on a 200-kb region (chr11:61,500,000-61,700,000), covering all three *FADS* genes. Variants were included in the analysis if they fulfilled the following criteria: 1) biallelic; 2) minor allele frequency (MAF) in the sample not less than 5%; 3) with rsID; 4) *p* value for Hardy-Weinberg equilibrium test larger than 0.001. Analysis was performed separately for the combined UK10K cohort and each of the five European populations in 1000G.

Haplotype network analysis was performed with an R software package, pegas^76^. To reduce the number of SNPs and thus the number of haplotypes included in the analysis, we restricted this analysis to part of the 85 kb *FADS1-FADS2* LD block, starting 5 kb downstream of *FDAS1* to the end of the LD block (a 60-kb region). To further reduce the number of SNPs, in the analysis with all 1000GP European samples, we applied an iterative algorithm^77^ to merge haplotypes that have no more than three nucleotide differences by removing the differing SNPs. The algorithm stops when all remaining haplotypes are more than 3 nucleotides away. With this procedure, we were able to reduce the number of total haplotypes from 81 to 12, with the number of SNPs decreased from 88 to 34 (Supplementary Fig. S30). This set of 34 representative SNPs was used in all haplotype-based analysis in aDNA, 1000GP, HGDP and POPRES. Missing data (e.g. from a low imputation genotype probability) were included in the haplotype network analysis.

Of note, for the 12 haplotypes identified in 1000GP European samples, only five of them have frequency higher than 1% (Supplementary Table S2). These five haplotypes were designated as D, M1, M2, M3 and M4, from the most common to the least.

### Ancient DNA-based test for recent selection in Europe

The test was performed as described before^11^. Briefly, most European populations could be modelled as a mixture of three ancient source populations at fixed proportions. The three ancient source populations are West or Scandinavian hunter-gatherers (WSHG), early European farmers (EF), and Steppe-Ancestry pastoralist (SA) (Supplementary Table S1). For modern European populations in 1000G, the proportions of these three ancestral sources estimated at genome-wide level are (0.196, 0.257, 0.547) for CEU, (0.362, 0.229, 0.409) for GBR, (0, 0.686, 0.314) for IBS, and (0, 0.645, 0.355) for TSI. FIN was not used because it does not fit this three-population model^11^. Under neutrality, the frequencies of a SNP (e.g. reference allele) in present-day European populations are expected to be the linear combination of its frequencies in the three ancient source populations. This serves as the null hypothesis: *p_mod_ = CP_anc_*, where *p_mod_* is the frequencies in A modern populations, *p_anc_* is the frequencies in B ancient source populations while *C* is an AxB matrix with each row representing the estimated ancestral proportions for one modern population. The alternative hypothesis is that *p_m0d_* is unconstrained by *p*_*anc*_. The frequency in each population is modelled with binomial distribution: *L*(*p; D*) *= B*(*X, 2N, p*), where *X* is the number of designated allele observed while *N* is the sample size. In ancient populations, *X* is the expected number of designated allele observed, taking into account uncertainty in imputation. We write ℓ(*p; D*) for the log-likelihood. The log-likelihood for SNP frequencies in all three ancient populations and four modern populations are: 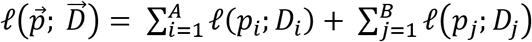. Under the null hypothesis, there are B parameters in the model, corresponding to the frequencies in B ancient populations. Under the alternative hypothesis, there are A+B parameters, corresponding to the frequencies in A modern populations and B ancestral populations. We numerically maximized the likelihood separately under each hypothesis and evaluate the statistic (twice the difference in log-likelihood) with the null *χ_A_^2^* distribution. Inflation was observed with this statistic in a previous genome-wide analysis and a *λ* = 1.38 was used for correction^11^. Following this, we applied the same factor in correcting the *p* values in our analysis. For genotyped SNPs previously tested, similar scales of statistical significance were observed as in the previous study (Supplementary Fig. 31). We note that for the purpose of refining the selection signal with imputed variants, only relative significance levels across variants are informative.

In addition to combining signals from four present-day European populations, we further performed tests separately in the two South European populations (IBS and TSI) and in the two North European populations (CEU and GBR). In these two cases, A = 2 and the null distribution is *χ*_2_^2^. For comparison between the North and the South, we used three statistics: the final *p* value, the maximum likelihood under the null hypothesis, and the maximum likelihood under the alternative hypothesis.

### Ancient DNA-based test for ancient selection in pre-Neolithic European hunter-gatherers

Two Bayesian methods, the Schraiber *et al.* method^24^ and the ApproxWF^25^, were applied to infer natural selection from allele frequency time series data. The two software were downloaded from https://github.com/Schraiber/selection and https://bitbucket.org/phaentu/approxwf/downloads/, respectively. The Schraiber *et al.* method models the evolutionary trajectory of an allele under a specified demographic history and estimates selection coefficients (*s*_1_ and *s*_2_) for heterozygotes and homozygotes of the allele under study. This method has two modes, with or without the simultaneous estimation of allele age. Without the estimation of allele age, this method models the frequency trajectory only between the first and last time points provided and its estimates of selection coefficients describe the selection force during this period only. With the simultaneous estimation of allele age, this method models the frequency trajectory starting from the first appearance of the allele to the last time point provided. In this case, the selection coefficients describe the selection force starting from the mutation of the allele, which therefore should be the derived allele. For demographic history, we used two models: a constant population size model with N_e_=10,000 and a more realistic model with two historic epochs of bottleneck and recent exponential growth^26^. However, the recent epoch of exponential growth does not have an impact on our analysis because for our analysis the most recent sample, WSHG, has an age estimate of ∼7500 years, predating the onset of exponential growth (3520 ya, assuming 25 years per generation). ApproxWF can simultaneously estimate selection coefficient and demographic history (only for constant population size model). For our purpose, we set the demographic history as N_e_=10,000. It estimates selection coefficient for homozygotes, *s*, and dominance coefficient, *h*. The selection coefficient estimated is for the time points specified by the input data.

Four groups of pre-Neolithic European hunter-gatherers were included in our test: the Věstonice cluster (median sample age: 30,076 yo), the El Mirón cluster (14,959 yo), the Villabruna cluster (10,059 yo) and WSHG (7,769 yo). To identify SNPs with evidence of positive selection during the historic period from Věstonice to WSHG, we applied both methods on most SNPs in the *FADS* locus. The Schraiber *et al.* method was run twice with two demographic models while ApproxWF was run once with the constant size model. For the two candidate SNPs (rs174570 and rs2851682), we further ran the Schraiber *et al.* method with the more realistic demographic model to simultaneously estimate their selection coefficients and allele ages. Statistical significance was considered if the 95% CI of selection coefficient does not overlap with 0. Details about running the two software were in Supplementary Methods.

### Modern DNA-based selection tests

We performed two types of selection tests for modern DNAs: site frequency spectrum (SFS)-based and haplotype-based tests. These tests were performed separately in each of the five European populations from 1000G and each of the two cohorts from UK10K. For SFS-based tests, we calculated genetic diversity (π), Tajima’s D^78^, and Fay and Wu’s H^79^, using in-house Perl scripts. We calculated these three statistics with a sliding-window approach (window size = 5 kb and moving step = 1 kb). Statistical significance for these statistics were assessed using the genome-wide empirical distribution. Haplotype-based tests, including iHS^80^ and nSL^21^, were calculated using software selscan (version 1.1.0a)^81^. Only common biallelic variants (MAF > 5%) were included in the analysis. Genetic variants without ancestral information were excluded. These two statistics were normalized in frequency bins (1% interval) and the statistical significance of the normalized iHS and nSL were evaluated with the empirical genome-wide distribution. The haplotype bifurcation diagrams and EHH decay plots were drawn using an R package, rehh^82^. Singleton Density Score (SDS) based on UK10K was directly retrieved from a previous study^22^.

### Geographical frequency distribution analysis

For plots of geographical frequency distribution, the geographical map was plotted with an R software package, maps (https://CRAN.R-project.org/package=maps) while the pie charts were added with the mapplots package (https://cran.r-project.org/web/packages/mapplots/index.html). Haplotype frequencies were calculated based on haplotype network analysis with pegas^76^, which groups haplotypes while taking into account missing data. SNP frequencies were either the observed frequency, if the SNP was genotyped, or the expected frequency based on genotype probability, if the SNP was imputed.

### Targeted association analysis for SNP rs174594 in UK10K

We performed association analysis for rs174594 in two UK10K datasets - ALSPAC and TwinsUK^72^. For both datasets, we analyzed height, weight, BMI and lipid-related traits including total cholesterol, low density lipoprotein, very low density lipoprotein, high density lipoprotein, Apolipoprotein A-I (APOA1), Apolipoprotein B (APOB) and triglyceride. We performed principal components analysis using smartpca from EIGENSTRAT software^83^ with genome-wide autosomal SNPs and we added top 4 principal components as covariates for all association analysis. We also used age as a covariate for all association analysis. Sex was added as a covariate only for ALSPAC dataset since all individuals in TwinsUK dataset are female. For all lipid-related traits, we also added BMI as a covariate.

## Data availability

Ancient DNA: https://reich.hms.harvard.edu/datasets

1000 Genomes Project: ftp://ftp.1000genomes.ebi.ac.uk/vol1/ftp/release/20130502/

Human Genome Diversity Project (HGDP): http://www.hagsc.org/hgdp/files.html

Population Reference Sample (POPRES): dbGaP Study Accession: phs000145.v4.p2

UK10K: https://www.uk10k.org/data_access.html

Singleton Density Score (SDS): https://github.com/yairf/SDS

## Code availability

Most analyses were conducted with available software and packages as described in the respective subsections of Methods. Customized Perl and R scripts were used in performing site frequency spectrum-based selection tests, and for general plotting purposes. All these scripts are available upon request.

## Acknowledgements

We thank Montgomery Slatkin and Joshua Schraiber for their help in running their software, David Reich and Iain Mathieson for making their data publicly available, Leonardo Arbiza, Charles Liang, Daniel (Alex) Marburgh, Edward Li, Kumar Kothapalli, Tom Brenna, and all members of the Keinan lab for helpful discussion and comments on the manuscript. This work was supported by the National Institutes of Health (Grants R01HG006849 and R01GM108805 to AK) and the Edward Mallinckrodt, Jr. Foundation (AK).

This study makes use of data generated by the UK10K Consortium, derived from samples from UK10K_COHORT_ALSPAC and UK10K_COHORT_TWINSUK. A full list of the investigators who contributed to the generation of the data is available from www.UK10K.org. Funding for UK10K was provided by the Wellcome Trust under award WT091310.

The collections and methods for the Population Reference Sample (POPRES) are described by Nelson *et al.* (2008). The datasets used for the analyses described in this manuscript were obtained from dbGaP at http://www.ncbi.nlm.nih.gov/projects/gap/cgi-bin/study.cgi?study_id=phs000145.v1.p1 through dbGaP accession number phs000145.v1.p1.

## Author contributions

A.K. and K.Y. conceived and designed the project. K.Y. performed data collection and analysis, with contributions from D.W. and F.G.. K.Y. and A.K. interpreted the results, with contribution from O.B. on the anthropological perspective. K.Y. and A.K. wrote the manuscript. All authors read, edited and approved the final version of the manuscript.

## Competing interests

The authors declare no competing interests.

